# A translational MRI approach to validate acute axonal damage detection

**DOI:** 10.1101/2022.04.27.489694

**Authors:** Nicola Toschi, Antonio Cerdán Cerdá, Constantina A Treaba, Valeria Barletta, Elena Herranz, Ambica Mehndiratta, Caterina Mainero, Silvia De Santis

**Author notes:** **Corresponding Author:** Silvia De Santis, Instituto de Neurociencias, avda Ramon y Cajal S/N, 03550 San Juan de Alicante (Spain).

## Abstract

Axonal degeneration is a central pathological feature of neurodegenerative pathologies and is closely associated with irreversible clinical disability. Current noninvasive methods to detect axonal damage *in vivo* are limited in their specificity, clinical applicability, and lack of proper validation. We aimed to validate an MRI framework based on multicompartment modeling of the diffusion-weighted signal (AxCaliber) in rats in the presence of axonal pathology, achieved through injection of a neurotoxin damaging the neuronal terminal of axons. We then applied the same MRI protocol to map axonal integrity in the whole brain of multiple sclerosis relapsing-remitting patients and age-matched healthy controls, a pathology associated with a neurodegenerative component.

AxCaliber is sensitive to microstructural axonal damage in rats, as demonstrated by a significant increase in the mean axonal caliber along the target tract, which correlated with the neurotoxin neurofilament staining. In humans, we uncovered a diffuse increase in mean axonal caliber in multiple sclerosis lesions and, importantly, in most areas of the normal-appearing white matter. Our results demonstrate that axonal diameter mapping is a sensitive and specific imaging biomarker able to link noninvasive imaging contrasts with the underlying biological substrate, supporting the key role of generalized axonal damage useful in diseases such as multiple sclerosis.

## Introduction

Magnetic resonance imaging (MRI), and particularly diffusion-based approaches, have been applied to investigate axonal damage in white matter (WM) (Inglese and Bester, 2010). However, conventional diffusion MRI is notoriously nonspecific to different tissue compartments, such as axons or myelin (De Santis et al., 2014), hampering the ability to distinguish axonal damage from other microstructural changes. In addition, due to the limitations of quantitatively comparing *in vivo* to *ex vivo* data, validation of imaging findings is rarely performed (Horowitz et al., 2015b).

Axonal damage is a main pathological substrate of irreversible neurological disability in multiple sclerosis (MS). In MS, axonal damage can either be direct or secondary to demyelination, glial activation or exposure to excitatory amino acids and cytokines (Haines et al., 2011). As the transition to progressive MS occurs when an axonal loss threshold is reached and the brain compensatory capacity is surpassed (Criste et al., 2014), the development of novel in vivo, noninvasive strategies for characterizing axonal microstructure becomes essential for early disease detection. While animal models recapitulating MS pathophysiology are available (Torkildsen et al., 2008),(Constantinescu et al., 2011), none of these focus on axonal damage.

AxCaliber is a novel imaging framework able to estimate axonal diameter (Assaf et al., 2008), which takes advantage of recent hardware advances (Jones et al., 2018) combined with advanced multicompartmental modeling of the diffusion MRI signal. AxCaliber has revealed higher axonal diameter in the normal-appearing WM (NAWM) of the corpus callosum of MS patients than in healthy controls, which was interpreted as a sign of axonal damage (Huang et al., 2016). However, in its original formulation, AxCaliber is only applicable to voxels characterized by a single fiber orientation (such as the corpus callosum (Huang et al., 2016)), while at least 70% of the brain voxels contain two or more dominant fiber orientations (Jeurissen et al., 2013). Recently, we used a modified AxCaliber model to map whole-brain axonal diameter (De Santis et al., 2019b), hence providing a means to characterize axonal damage in a whole-brain fashion.

From a pathophysiological point of view, measuring a higher estimated axonal diameter does not, *per se*, represent conclusive evidence of axonal damage. Indeed, MRI sensitivity to the loss axonal loss of axons has yet to be fully demonstrated to establish its clinical utility. Recently, we proposed an approach to selectively manipulate specific microstructural aspects of the parenchyma through targeted, unilateral injection of neurotoxic agents (Garcia-Hernandez et al., 2022).

Here, we aimed to characterize axonal damage using a translational approach. Our specific objectives were i) to validate AxCaliber-based axonal mapping in a preclinical model of fimbria damage induced by stereotaxic injection of ibotenic acid into the hippocampus and ii) to employ the human AxCaliber MRI protocol to investigate changes in axonal microstructure in WM lesions and NAWM in MS brains. Specifically, we iia) compared whole-brain, voxelwise axonal diameter in MS patients relative to healthy controls iib) compared the axonal diameter in lesions to that in contralateral NAWM in MS patients as well as in the corresponding location in a cohort of healthy controls; and iic) investigated differences in axonal diameter between lesions classified as isointense or hypointense on T1-weighted scans. The latter investigation was motivated by the proposed relationship between the intensity of lesions in T1 weighted imaging and the degree of relative preservation versus more severe damage of the myelin/axonal structure (Loevner et al., 1995).

## Materials and Methods

### Animal preparation

All animal experiments were approved by the Institutional Animal Care and Use Committee of the Instituto de Neurociencias de Alicante, Alicante, Spain, and comply with the Spanish (law 32/2007) and European regulations (EU directive 86/609, EU decree 2001-486, and EU recommendation 2007/526/EC; Project MRI-STRUCTURE 677/2018). The ARRIVE 10 checklist was used. Animal preparation (n=9 rodents) was carried out as described here (Garcia-Hernandez et al., 2022). Briefly, axonal damage in the fimbria was achieved by injecting 1 μl of saline and ibotenic acid (a selective agonist of N-methyl-D-aspartate (NMDA) glutamate receptors that produces selective neurotoxicity(Zinkand et al., 1992)) at a concentration of 2.5 μg/μl in the right dorsal hippocampus (coordinates bregma −3.8 mm, sup-inf 3.0 mm, 2 mm from midline in the left hemisphere). The opposite hemisphere was injected with saline only so that each animal had its own control (Fig. 1a). Neuronal degeneration in the hippocampus translates into axonal loss in its major axonal output bundle, the fimbria, which is therefore used as a model for Wallerian-like axonal degeneration (Conforti et al., 2014). Fourteen days after surgery, rats underwent MRI scans using the AxCaliber protocol and were immediately perfused for *ex vivo* MRI immunohistological analysis. Histological analysis was used to stain neuronal somas (NeuN) and quantify neuronal death in the hippocampus, neurofilaments and myelin basic protein (MBP) to quantify axonal integrity and myelination in the fimbria, respectively.

**Figure 1:**
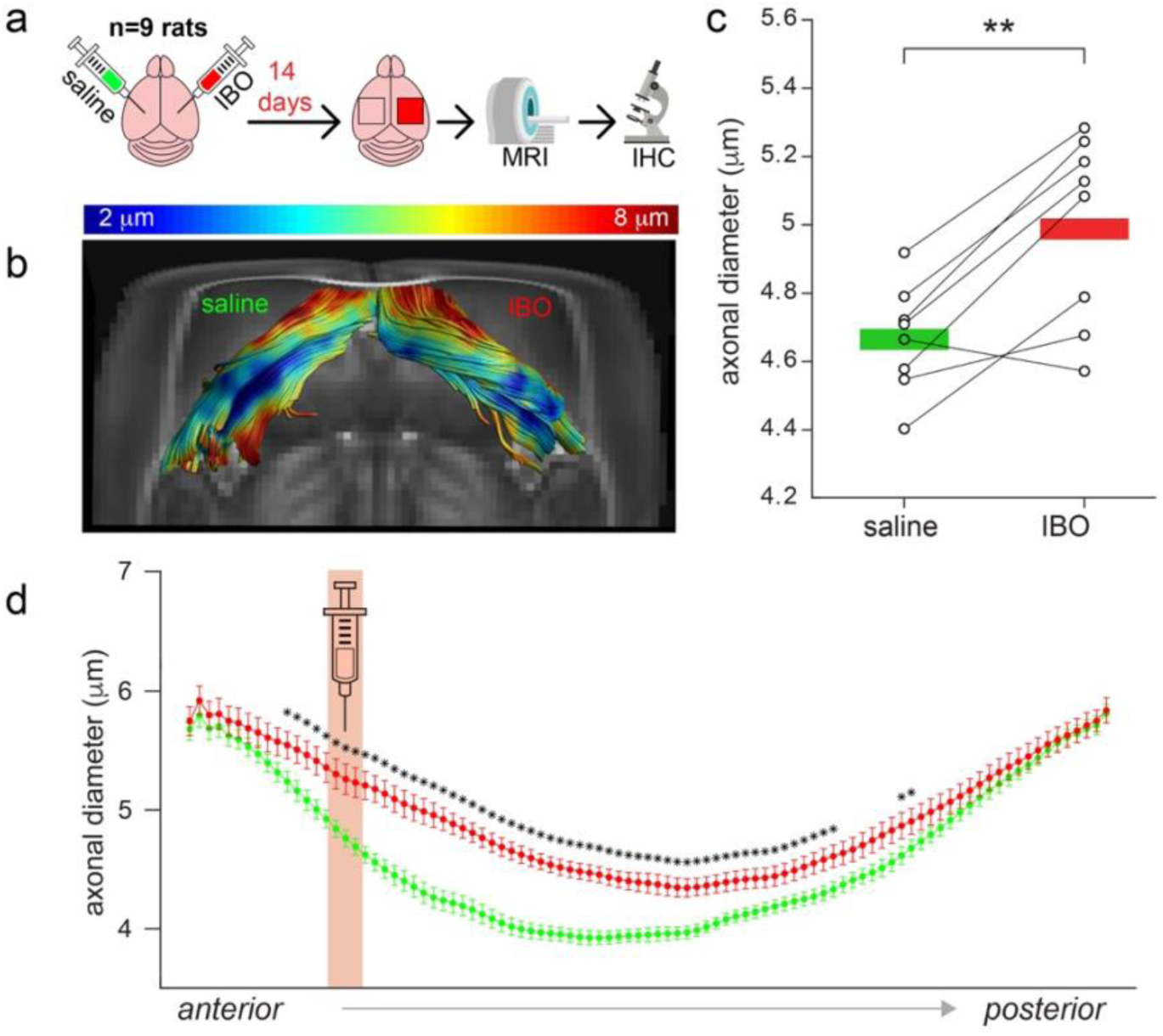
Experimental model of axonal damage. a) Experimental scheme of stereotaxic injections of ibotenic acid (IBO) in the left hippocampus of n=9 rats. The right hippocampus was injected with saline solution and used as a control. b) An example of the tractography of the fimbria from one representative animal. Axonal diameter is projected on the tract in the form of color coding. c) Mean axonal diameter calculated in the ibotenic vs saline-injected fimbria reconstructed using tractography. Asterisks represent significant differences (paired t test across hemispheres, p<0.01). d) Mean and standard deviation across subjects of axonal diameter measured across all the streamlines constituting the fimbria in the antero-posterior axis, divided into 100 steps. The injection site is shown in red. Asterisks represent significant differences between injected and control fimbrias in post hoc t tests, corrected for multiple comparisons across tract locations.

### Subjects

The local institutional review board approved this study, and written informed consent was obtained from all participants. Ten MS patients (age range 27-59, 7 females) and 6 controls (age range 23-53, 3 females) participated in the study. The sample size was calculated based on previous literature (Huang et al., 2016). Age was not significantly different between the two groups (p=0.093, Mann–Whitney U Test). Eligibility criteria in patients were a diagnosis of relapsing-remitting MS (Polman et al., 2011), being on stable disease-modifying treatment or no treatment for at least 3 months, absence of clinical relapse within 3 months and absence of corticosteroid use within one month from study enrollment. Physical disability was assessed by a neurologist according to the Expanded Disability Status Scale (EDSS (Kurtzke, 1983)), and cognitive ability was assessed using the Symbol Digit Modalities Test (SDMT). Demographic and clinical data are shown in Table 1.

**Table 1.**
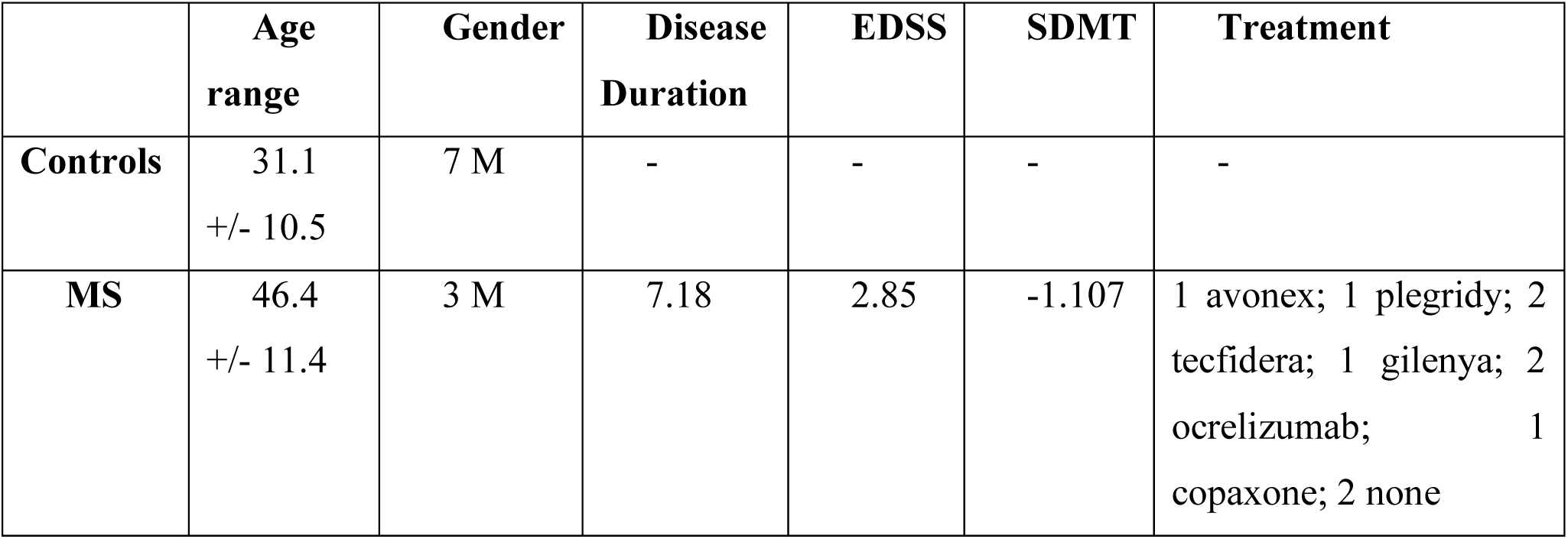
Demographic characteristics of the studied cohort, including age/gender, disease duration, EDSS, SDMT and MS treatment

### MRI acquisition

#### Rats

MRI was performed in vivo on a 7 T scanner (Bruker, BioSpect 70/30, Ettlingen, Germany). Dw-MRI data were acquired using an Echo Planar Imaging stimulated echo diffusion sequence, with 31 uniform distributed gradient directions, b=0(1), 2000(15) and 4000(15) s/mm^2^, diffusion times (Δ) 15, 25, 40 and 60 ms, repetition time (TR) = 5000 ms and echo time (TE) = 25 ms, for a total of 124 volumes acquired. Fourteen slices were set up centered in the fimbria with field of view (FOV) = 25×25 mm^2^, matrix size = 110 × 110, in-plane resolution = 0.225×0.225 mm^2^ and slice thickness = 0.6 mm. The total acquisition time was 36 minutes.

#### Humans

All participants were scanned on a Siemens 3T Connectom scanner, a customized 3T MAGNETOM Skyra system (Siemens Healthcare, Erlangen, Germany) housed at the MGH/HST Athinoula A. Martinos Center for Biomedical Imaging, Boston, Massachusetts, USA. The Connectom scanner is equipped with gradient coils capable of generating 300 mT/m maximum gradient strength, hence allowing minimization of δ (gradient duration) and echo times even at high b-values. A 64-channel brain array coil(Keil et al., 2013) was used for data acquisition. Dw-MRI data were acquired using an Echo Planar Imaging diffusion sequence, with 91 uniformly distributed gradient directions, b=0(1), 2000(30) and 4000(60) s/mm^2^, diffusion times (Δ) 17, 35 and 61 ms with four non-diffusion weighted images, TR = 5000 ms and TE = 89 ms, for a total of 273 volumes acquired. Eighty-two slices were set up to cover the whole brain with FOV = 220×220 mm^2^, matrix size = 110×110, in-plane resolution = 2×2 mm^2^ and slice thickness = 2 mm. In addition, anatomical images were acquired using 3D sequences with a 1.0 mm isotropic voxel size: T_1_-weighted multiecho magnetization-prepared rapid gradient-echo images were acquired in all participants(van der Kouwe et al., 2008). Fluid-attenuation inversion recovery (FLAIR) images were also acquired in MS patients for white matter lesion segmentation. The total acquisition time was 24 minutes.

### Tissue processing and immunohistochemistry

Rats were deeply anesthetized with a lethal dose of sodium pentobarbital, 46 mg/kg, injected intraperitoneally (Dolethal, E.V.S.A. laboratories., Madrid, España). Rats were then perfused intracardially with 100 ml of 0.9% phosphate saline buffer (PBS) and 100 ml of ice-cold 4% paraformaldehyde (PFA, BDH, Prolabo, VWR International, Louvain, Belgium). Then, brains were immediately extracted from the skull and fixed for 1 hour in 4% PFA. Afterwards, brains were included in 3% Agarose/PBS (Sigma-Aldrich, Madrid, Spain), and cut in vibratome (VT 1000S, Leica, Wetzlar, Germany) in 50 μm thick serial coronal sections.

Coronal sections were rinsed and permeabilized three times in 1xPBS with Triton X-100 at 0.5% (Sigma-Aldrich, Madrid, Spain) for 10 minutes each and then blocked in the same solution with 4% of bovine serum albumin (Sigma-Aldrich, Madrid, Spain) and 2% of goat serum donor herd (Sigma-Aldrich, Madrid, Spain) for 2 hours at room temperature. The slices were then incubated overnight at 4ºC with primary antibodies for myelin basic protein (1:250 Millipore Cat# MAB384-1ML, RRID:AB_240837), neurofilament 160 kD medium (1:250, Abcam Cat# ab134458, RRID:AB_2860025) and NeuN (1:250, Millipore Cat# MAB377, RRID:AB_2298772) to label myelin, axonal processes and nuclei, respectively. The sections were subsequently incubated in specific secondary antibodies conjugated to the fluorescent probes, each at 1:500 (Molecular Probes Cat# A-11029, RRID:AB_2534088; Molecular Probes Cat# A-11042, RRID:AB_2534099) for 2h at room temperature. Sections were then treated with 4′,6-Diamidine-2′-phenylindole dihydrochloride at 15mM (DAPI, Sigma-Aldrich, Madrid, Spain) during 15 minutes at room temperature. Finally, sections were mounted on slides and covered with an anti-fading medium using a mix solution 1:10 Propyl-gallate:Mowiol (P3130, SIGMA-Aldrich, Madrid, Spain; 475904, MERCK-Millipore, Massachussets, United States). For myelin labelling, antigen retrieval was performed in 1% citrate buffer (Sigma-Aldrich, Madrid, Spain) and 0.05% of Tween 20 (Sigma-Aldrich, Madrid, Spain) warmed to 80ºC for protein unmasking.

The tissue sections were then examined using a computer-assisted morphometry system consisting of a Leica DM4000 fluoresce microscope equipped with a QICAM Qimaging camera 22577 (Biocompare, San Francisco, USA) and Neurolucida morphometric software (MBF, Biosciences, VT, USA). Myelin, neurofilament and neural nuclei fluorescent analysis was performed using Icy software (de Chaumont et al., 2012). For neural nuclei, two ROIs of 200 µm^2^ were placed per hippocampus per hemisphere in at least 5 slices per rat to obtain the corresponding intensity values. Similarly, for MBP and neurofilaments, a ROI of 400 µm^2^ were placed per fimbria per hemisphere in at least 5 slices per rat.

### Data analysis

Paired t tests were used to assess the differences in histological quantities between injected versus control hemispheres. One animal did not present sign of neuronal damage in the hippocampus, and was therefore excluded from the analysis.

Diffusion-weighted rat data were preprocessed as described here (De Santis et al., 2019a). Human diffusion-weighted data were preprocessed with software tools in FreeSurfer V5.3.0 and FSL V5.0. Preprocessing included gradient nonlinearity correction, motion correction and eddy current correction, including corresponding b-matrix reorientation. Additional preprocessing details are available at http://www.humanconnectome.org/. In both rats and humans, MRI data were employed to fit the AxCaliber model using in-house software written in MATLAB R2015b (The Mathworks) to extract the average axonal diameter, as described here (De Santis et al., 2019b). Importantly, the model includes fiber dispersion by accounting for more than a single fiber orientation.

For all MS patients, lesion masks were segmented on the FLAIR images using a semiautomated method (3D-Slicer v4.2.0), and contralateral NAWM were created through coregistration. The intensity of the lesions in the T1-weighted image was used to classify lesions into hypo- and isointense (van den Elskamp et al., 2008).

In rats, the fimbria was reconstructed bilaterally using the DTI-based tractography (Fig. 1b) algorithm in the software ExploreDTI (Alexander Leemans et al., n.d.), which was set to employ the lowest Δ and b-value. Mean axonal diameter values along the tracts were obtained for each animal in both hemispheres. Paired t tests were used to assess differences in estimates of the axonal diameter between the injected and contralateral fimbria, and repeated-measures ANOVA followed by correction for multiple comparisons across tract locations (factors: tract location, treatment (ibotenic acid vs saline) and tract location*treatment) was used to further analyze the effect of ibotenic acid injection as a function of distance from the injection site.

For groupwise analysis of NAWM, we employed a previously detailed approach (De Santis et al., 2019a). Briefly, FA maps (calculated using the lowest Δ and b-value) were employed to initialize the first steps of an improved version of the TBSS (Smith et al., 2006). In this version, the coregistration steps are performed using extremely accurate tools (Klein et al., 2009, p. 14). The warping procedure accounts for lesion masks by excluding them from the similarity metric calculation of a permutation-based nonparametric inference approach to general linear modeling (Winkler et al., 2014). We tested for significant differences in the estimated axonal diameter within NAWM of MS patients compared to controls while excluding lesion masks and controlling for age and multiple comparisons across clusters using threshold-free cluster enhancement. In addition, we also tested for voxelwise associations of estimates of axonal diameter with EDSS, SDMT and disease duration while correcting for age and multiple comparisons across clusters.

To compare the lesions with topographically homologous regions, the corresponding nonlesioned contralateral region was segmented by applying a nonlinear transform mapping the 3D T1-weighted image to its corresponding right-left flipped mirror image. This was followed by visual inspection of the images to exclude possible lesioned tissue in the contralateral mask; voxels whose contralateral mask also contained lesioned tissue were excluded. Lesion masks were further refined by including only voxels that overlapped with each subject’s WM skeleton (as generated through the TBSS procedure) in native space to reduce the contribution of voxels with >2 main fiber orientations. For each MS patient, mean values for the axonal diameter were calculated in native space both within each lesion and within the corresponding contralateral NAWM region. Given that different anatomical locations present different baseline values for each parameter, we used a normalized summary statistic that eliminated this effect (normalized index of asymmetry (NIA), a normalized metric that quantifies relative discrepancies between homologous quantities while eliminating variability due to anatomical location), as defined in (De Santis et al., 2019a). Last, to compare values of the axonal diameter in lesions, NAWM and control WM, for each MS patient, we computed the average value of axonal diameter in scans from healthy controls within ROIS defined by the lesion mask after mapping the lesion masks onto this population. For this analysis, lesions were manually classified by an experienced operator as isointense or hypointense based on the contrast on the T1-weighted scan. Groups were compared using a general linear model (GLM) with lesion type (T1 iso and hypointense) as the “between” factor and cohort (healthy and MS) as the “within” factor. The model also included a group*lesion type interaction term and was followed by post hoc t tests corrected for multiple comparisons whenever a significant effect was detected.

## Results

### A rat model of acute axonal damage

Fourteen days after injection of the neurotoxin ibotenic acid into the hippocampus, DTI-based tractography reconstructions revealed a significant increase in the mean estimated axonal diameter (p=0.003) in the fimbria belonging to the injected hemisphere compared to the control (Fig. 1c), confirming that the damage affected a large portion of the tract. We revealed a significant effect of the injection (F_1,7_=19.5, p=0.003), of the position along the tract (F_98,686_=219.5, p<0.001) and of the interaction between them (F_98,686_=11.2, p=0.012); significant differences in mean estimated axonal diameter between injected and control tracts were mostly localized posterior to the injection site (Fig. 1d).

When comparing injected versus control hemispheres, we confirmed both neuronal loss in the hippocampus (p=0.026, Fig. 2a-b) and axonal damage in the fimbria (p=0.047, Fig. 2c-d), corresponding to a lower staining intensity of NeuN and higher intensity of neurofilament staining in the hemisphere injected with ibotenic acid. No differences were found in myelin content using MBP staining (Fig. S1), suggesting that at the studied time point, axonal structure was significantly altered, but the myelin sheath was still preserved. Neurofilament fluorescence intensity was significantly correlated with estimates of axonal diameter measured with MRI (r=0.54, p=0.029) in both injected and control hemispheres (Fig. 3).

**Figure 2:**
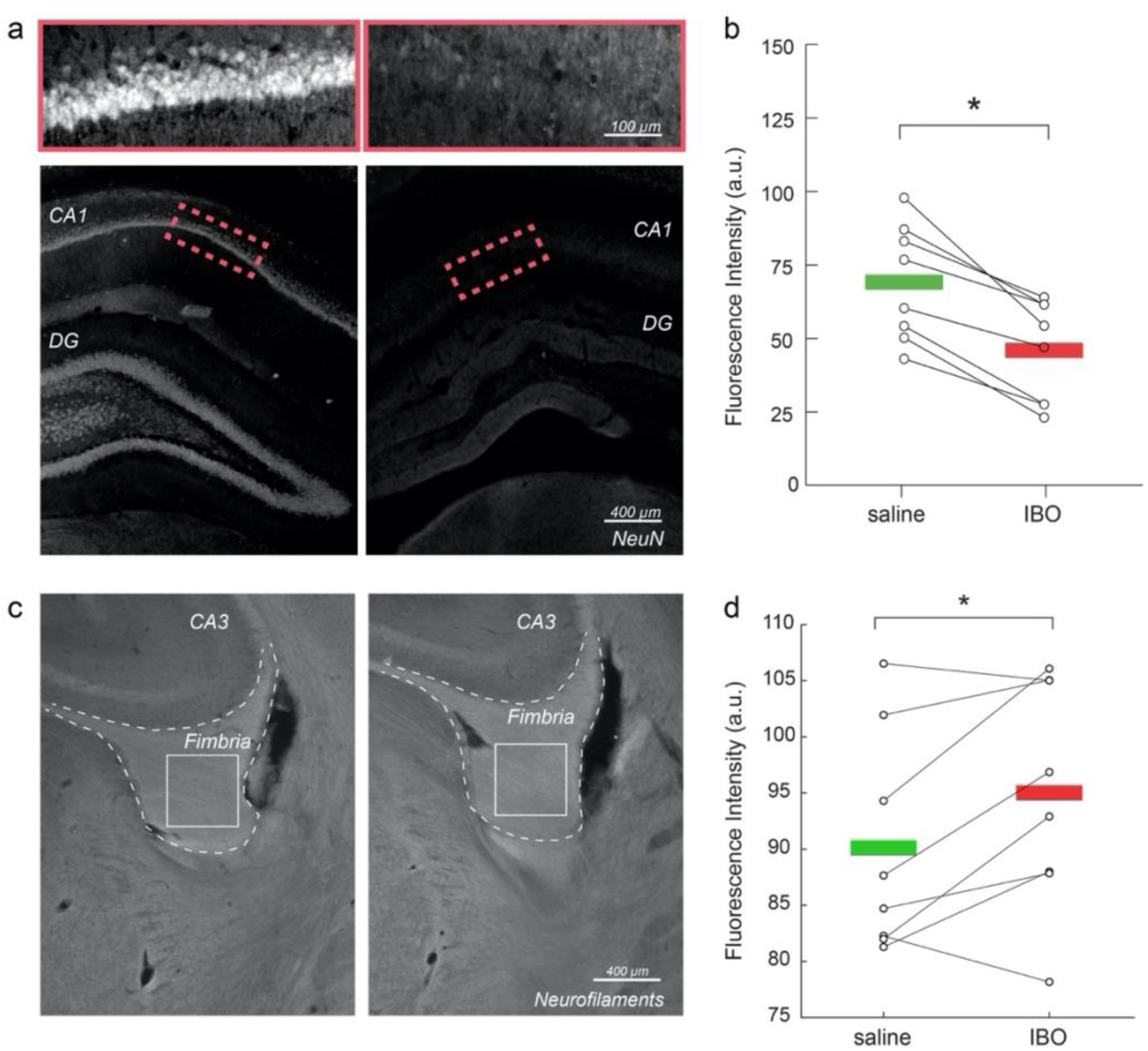
Immunohistochemical validation of axonal damage. a) NeuN staining in control vs. injected hippocampi. b) Mean NeuN intensity in control vs. injected hippocampi. Asterisks represent significant differences across hemispheres (paired t test, p<0.05) c) Neurofilament staining in control vs. injected fimbria. d) Mean neurofilament intensity in control vs. injected hippocampi. Asterisks represent significant differences in means across hemispheres (paired t test, p<0.05).

**Figure 3:**
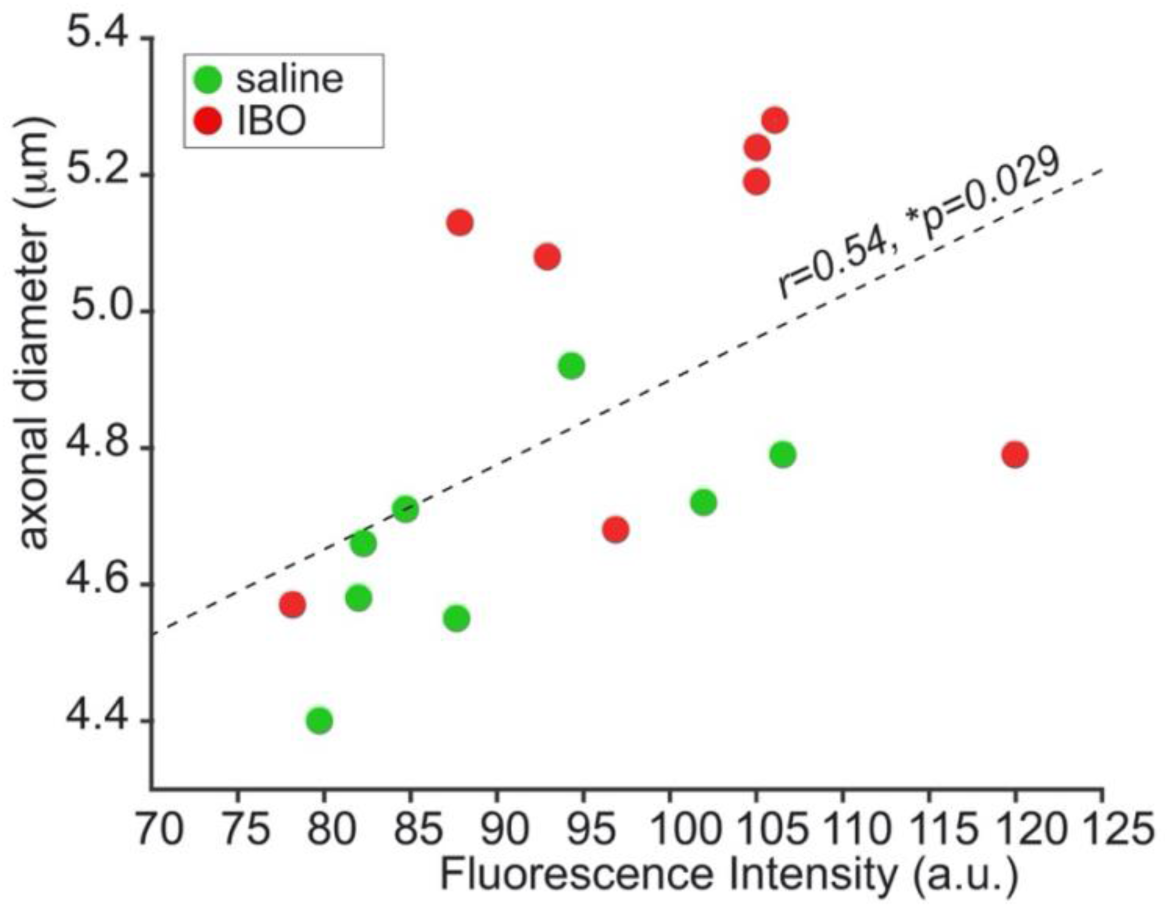
Correlation between MRI and histology. Significant correlation between neurofilament fluorescence intensity and axonal diameter measured with the Axcaliber model for all (ibotenic acid (red) and saline (green)) fimbrias.

### Axonal damage in normal-appearing white matter of multiple sclerosis patients

After preclinical validation in rats, we applied the clinical version of the AxCaliber MRI protocol to a cohort of MS patients and age-matched healthy controls. When comparing the axonal diameter in the NAWM of MS patients and controls, we found higher axonal diameter in the NAWM of the MS group (Fig. 4). The differences were mostly symmetrical across hemispheres and involved all major WM tracts, notably the corpus callosum, the internal capsule, the corona radiata, the thalamic radiation, the inferior longitudinal fasciculus, the cingulum, the fornix, the superior longitudinal fasciculus, the inferior fronto-occipital fasciculus, the uncinate fasciculus and the tapetum.

**Figure 4:**
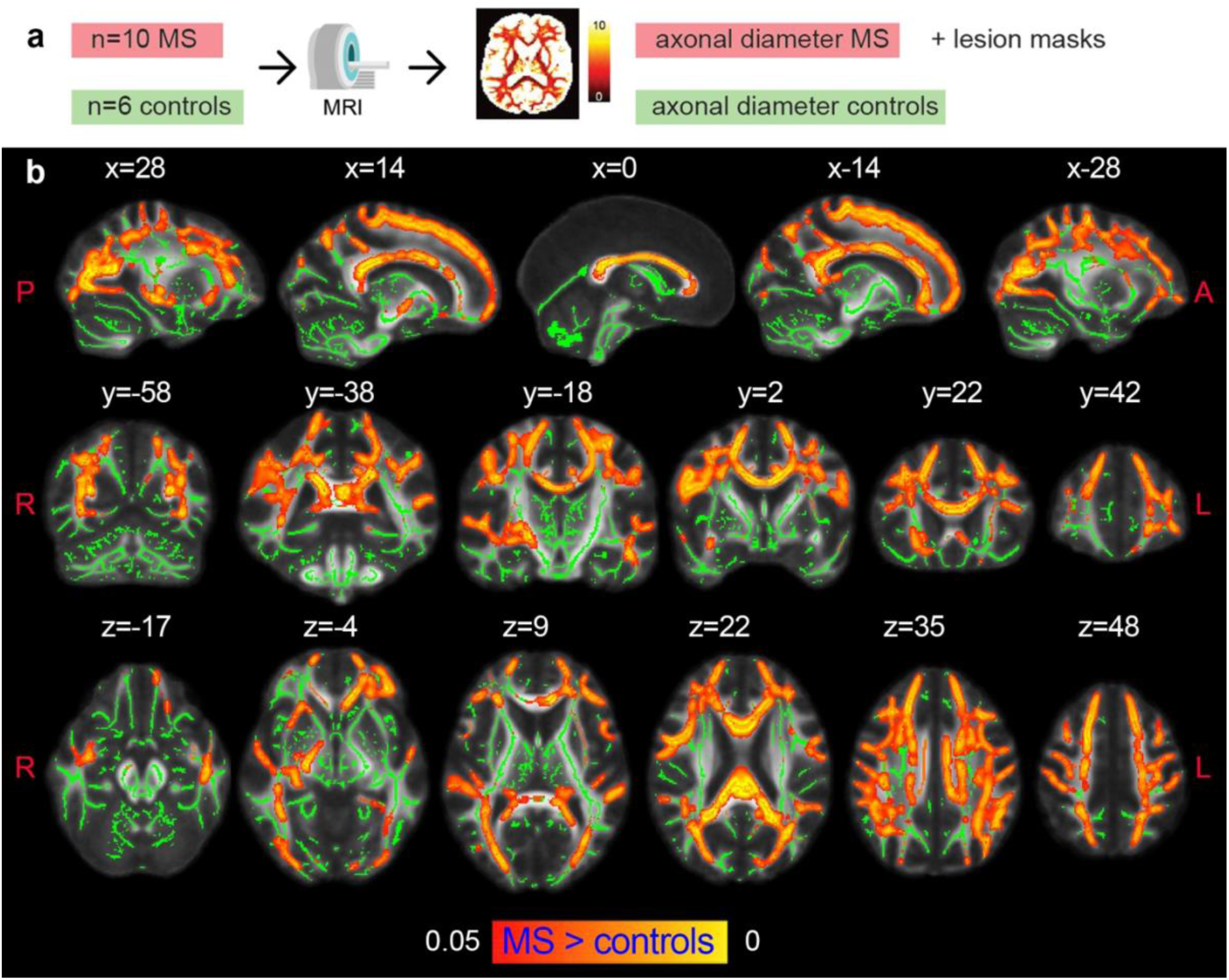
Axonal damage in MS normal-appearing white matter. a, Experimental scheme. b, Tract-based spatial statistics showing voxels in which the mean axonal diameter was significantly increased in multiple sclerosis versus control conditions (p<0.05, corrected). The opposite contrast was not statistically significant, so it is not shown in the figure. Green: skeletonized white matter. Red–Yellow: p value.

### Axonal pathology in multiple sclerosis lesions

Additionally, when examining MS lesions, we detected a larger axonal diameter in lesions compared to contralateral NAWM in MS patients, as indicated by a lower NIA in controls (Fig. 5c). When comparing axonal diameter in lesions with the average value in the corresponding regions defined across the healthy cohort, we found a significant group effect (F_1,235_=25.7, p<0.001) and of the interaction term group*lesion type (F_1,235_=40.1, p=0.03), after which we observed a selectively higher axonal diameter in T1 hypointense lesions only (Fig. 5 d-e).

**Figure 5:**
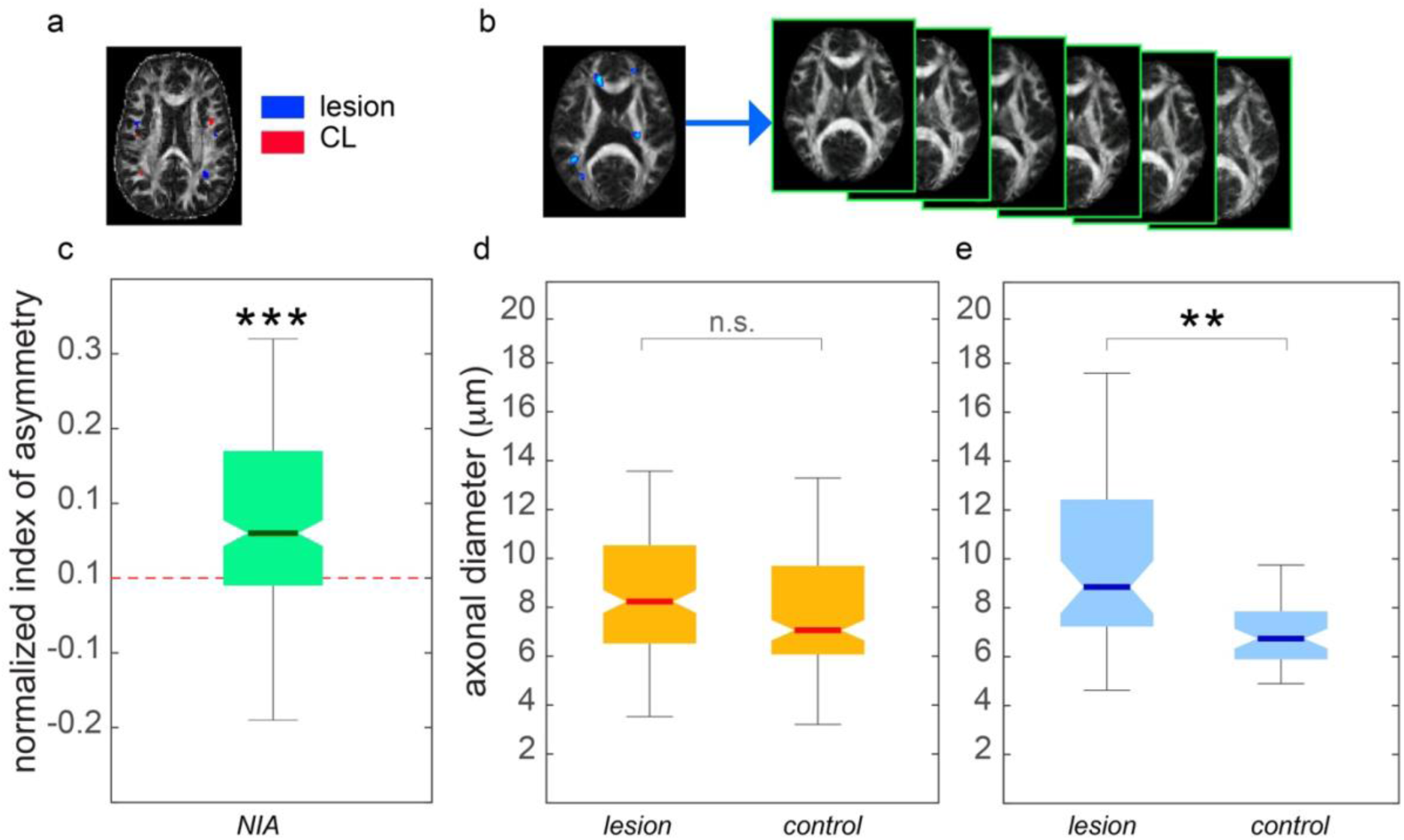
Axonal damage in MS lesions. a) Example of lesion and contralateral masks in one representative subject in native space, overlaid on a fractional anisotropy map. b) Example of a lesion mask in MNI space for one MS subject. The mask is used to calculate the mean axonal diameter across the healthy cohort in the anatomical region defined by the lesions. c) Normalized index of asymmetry (NIA) for mean axonal diameter between lesions and contralateral normal-appearing tissue. Asterisks represent statistically significant results from a one-sample t test (p<0.001), indicating that the NIA is significantly different from zero (in this case, positive), and hence, the axonal diameter in lesions is significantly larger than that in healthy tissue. d-e) Mean axonal diameter in lesions and in the same anatomical location as the lesions in healthy controls for isointense (d) and hypointense (e) lesions, according to T1-weighted contrast. ANOVA followed by post hoc comparisons highlighted significantly increased axonal diameter in lesions compared to controls in hypointense lesions only (p<0.01).

We did not find significant associations between estimates of axonal diameter and clinical variables (EDSS and SDMT) in either lesions or NAWM. However, we found a negative association at the trend level between estimates of axonal diameter and disease duration in the whole white matter skeleton in the TBSS analysis, which in ROI analysis was statistically significant in 2 ROIs (left cingulum and right inferior fronto-occipital fasciculus, see Fig. S2).

## Discussion

In this work, we used a preclinical model of acute axonal damage to demonstrate that MRI-based axonal diameter estimation is sensitive to early stages of axonal degeneration, even before alterations in the myelin sheet are detected. We then applied the same MRI preclinical protocol to a cohort of patients with MS and an age-matched healthy cohort, uncovering diffuse axonal damage in both the NAWM and in the lesioned WM tissue of MS patients.

Neuropathologically, axonal damage in MS manifests through the formation of varicosities and spheroids that enlarge the axonal diameter and are associated with impaired axonal transport (Criste et al., 2014). Accordingly, histological postmortem (Bergers et al., 2002) and animal studies (Nikić et al., 2011) report an increase in the mean axonal diameter in MS compared to control. This increase is also influenced by the higher vulnerability of smaller axons compared to larger ones (Tallantyre et al., 2010), which implies that smaller axons are lost earlier. Recently, histological alterations in axonal morphology have also been observed in NAWM in the absence of myelin abnormalities or inflammation (Luchicchi et al., 2021); this result could point to an imbalance of axon-myelin units as a primary event in MS pathogenesis, making noninvasive detection of axonal damage a priority.

Given the aforementioned neuropathological evidence, axonal damage in MS would reasonably manifest as an increase in axonal caliber measured with imaging. An increase in axonal diameter in MS was indeed reported in preliminary MRI studies(Huang et al., 2016), but the results were not supported by an independent validation. In the context of multicompartment models for diffusion signals, comparing imaging results with pathological evidence imaging results is fundamental to validating the model, as multicompartment models make numerous assumptions and simplifications (to cite a few: fixed diffusivities (Alexander et al., 2010), no exchange(Lasič et al., 2011), indirect account of the volume occupied by myelin (Assaf et al., 2008)).

Previous studies reported correlations between the axonal diameter measured with MRI and axonal caliber estimated using electron microscopy in healthy tissue (Barazany et al., 2009), but none thus far have demonstrated that AxCaliber is sensitive to axonal damage. Quantitative comparison between MRI maps and stained sections is severely hampered by the fixation process and other limitations (Horowitz et al., 2015b); here, we used a well-characterized rodent model in which the axonal compartment is selectively damaged. This approach has been previously used to prove the capability of diffusion MRI to dissect astrocyte and microglia activation in gray matter (Garcia-Hernandez et al., 2020). Here, we detected increased neurofilament staining intensity in the damaged tract, demonstrating altered axonal morphology, without alteration in the total amount of myelin. This change is picked up by MRI as an increase in the estimated axonal diameter. The significant correlation between neurofilament staining intensity and axonal diameter further validates the imaging parameter as a proxy for axonal damage. Overall, our preclinical results show that axonal diameter mapping through MRI can detect axonal pathology in vivo and can thus be used as a biomarker of axonal damage in MS and other neurological diseases.

In our work, we used an Axcaliber formulation that allows us to account for multiple fibers and thus access whole-brain axonal diameter mapping. Indeed, while previous findings focused on the corpus callosum(Huang et al., 2016), our results indicate an increase in mean axon diameter in MS patients in all the major tracts, demonstrating for the first time diffuse axonal pathology in NAWM. Whole-brain axonal diameter mapping also grants access to investigate axonal pathology in MS-related lesions. We found that the estimated axonal diameter is only significantly higher in T1-hypointense lesions, which are known to be characterized by a higher degree of axonal damage(Van Waesberghe et al., 1999). This suggests that axonal diameter mapping can be useful to classify lesions according to the severity of axonal pathology, which can be explored in the future as a new tool for patient stratification.

The lack of significant correlations with clinical variables (EDSS and SDMT) can possibly be explained by the relatively low sample size or potentially compensatory phenomena in our MS population, mainly composed of many early-stage patients.

This study has some limitations. The histological quantification of axonal damage is made using neurofilament staining and not electron microscopy, so we did not measure the axonal radius; however, neurofilaments are known to be altered in MS (Petzold, 2005). We preferred immunofluorescent staining over electron microscopy to correlate in vivo and ex vivo quantities due to the severe fixation process and the very small sample size needed for electron microscopy (Horowitz et al., 2015b).

As stated in the original paper, AxCaliber and related frameworks are somewhat less sensitive to small-diameter axons and therefore might overestimate the mean axon diameter. However, the fact that the estimated diameter, while possibly biased, captures meaningful variations across different geometries, with relevance to function, has been widely demonstrated (Horowitz et al., 2015a).

Despite some inevitable minor differences due to different brain sizes and magnet features, the human protocol was built to match the main characteristics of the preclinical diffusion sequence, such as the b-value and diffusion time range. Last, the method has been demonstrated preclinically only looking at one specific tract (the fimbria, as the tract with the largest number of hippocampal projections in the rat); however, it would be straightforward to extrapolate our results to other tracts. Future studies are needed to demonstrate this.

Despite the provided evidence in an animal model of axonal damage, we cannot exclude potential alternative explanations than an actual axonal diameter increase underlying the MS data, whose pathological substrate is inevitably more complex. In particular, MRI signals could include contributions from other cell populations, such as glia and oligodendrocytes. Further work is needed to account for all cell species in a more realistic model of WM.

In conclusion, given the central role of axonal pathology in MS, developing and validating strategies for early detection of axonal damage, in vivo and noninvasively, is of high priority; our results have the potential to improve early detection and monitoring of axonal pathology in the disease, as well as provide a novel imaging marker for monitoring the effects of treatments on the progression of axonal degeneration.

## Abbreviations

MS: Multiple Sclerosis;
NAWM: Normal Appearing White Matter;
MRI: Magnetic Resonance Imaging;
FLAIR: Fluid-attenuation inversion recovery;
EDSS: Disability Status Scale;
SDMT: Symbol Digit Modalities Test

## Data Availability

The preclinical data (imaging and histology) and the human imaging data that support the findings of this study will be made available upon acceptance for publication in the open repository Open Science Framework, and the references will be provided in the manuscript. The human data will be anonymized and skull-stripped prior to publication, according to the regulations of the institution where they were acquired (Massachusetts General Hospital, Boston, USA). The software used to process the imaging data is available in the open repository DIGITAL.CSIC (http://hdl.handle.net/10261/266923)

## Supplementary material

**Figure S1.**
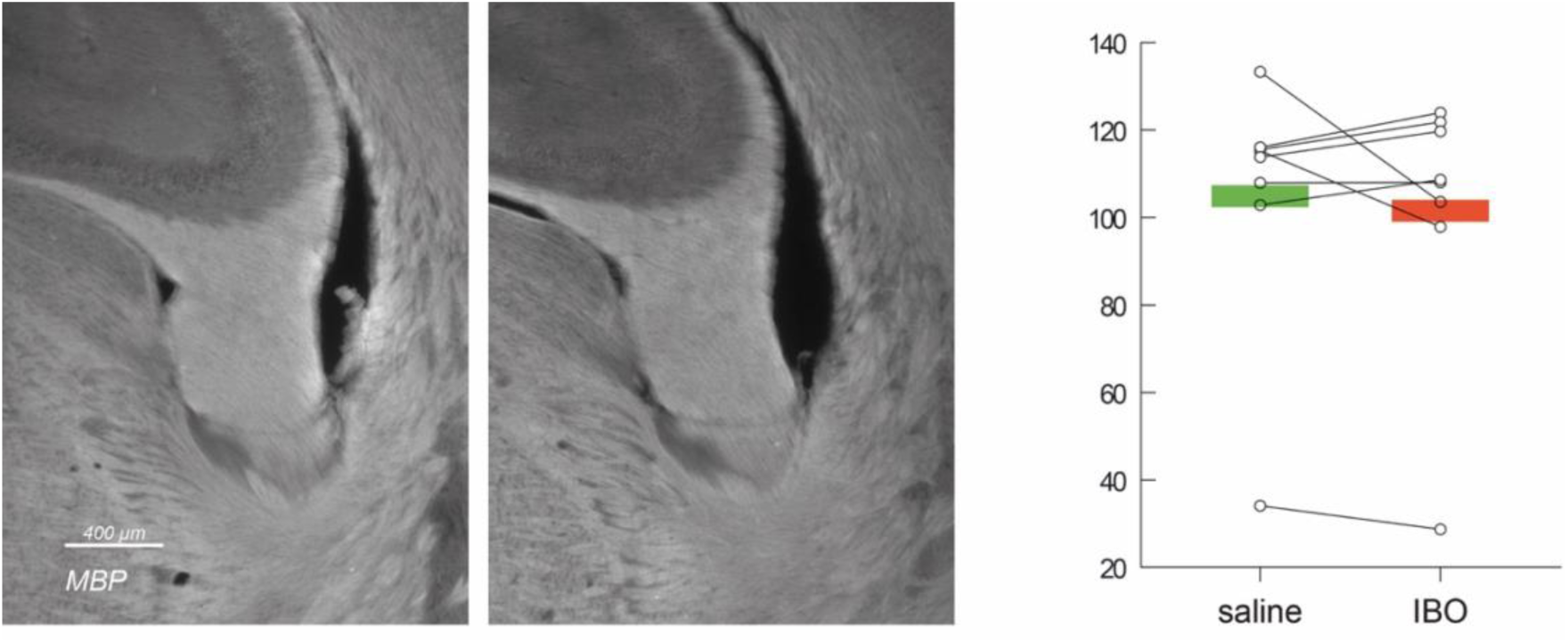
Myelin Basic Protein staining in injected versus control fimbria. No significant differences in myelination were found.

**Fig. S2.**
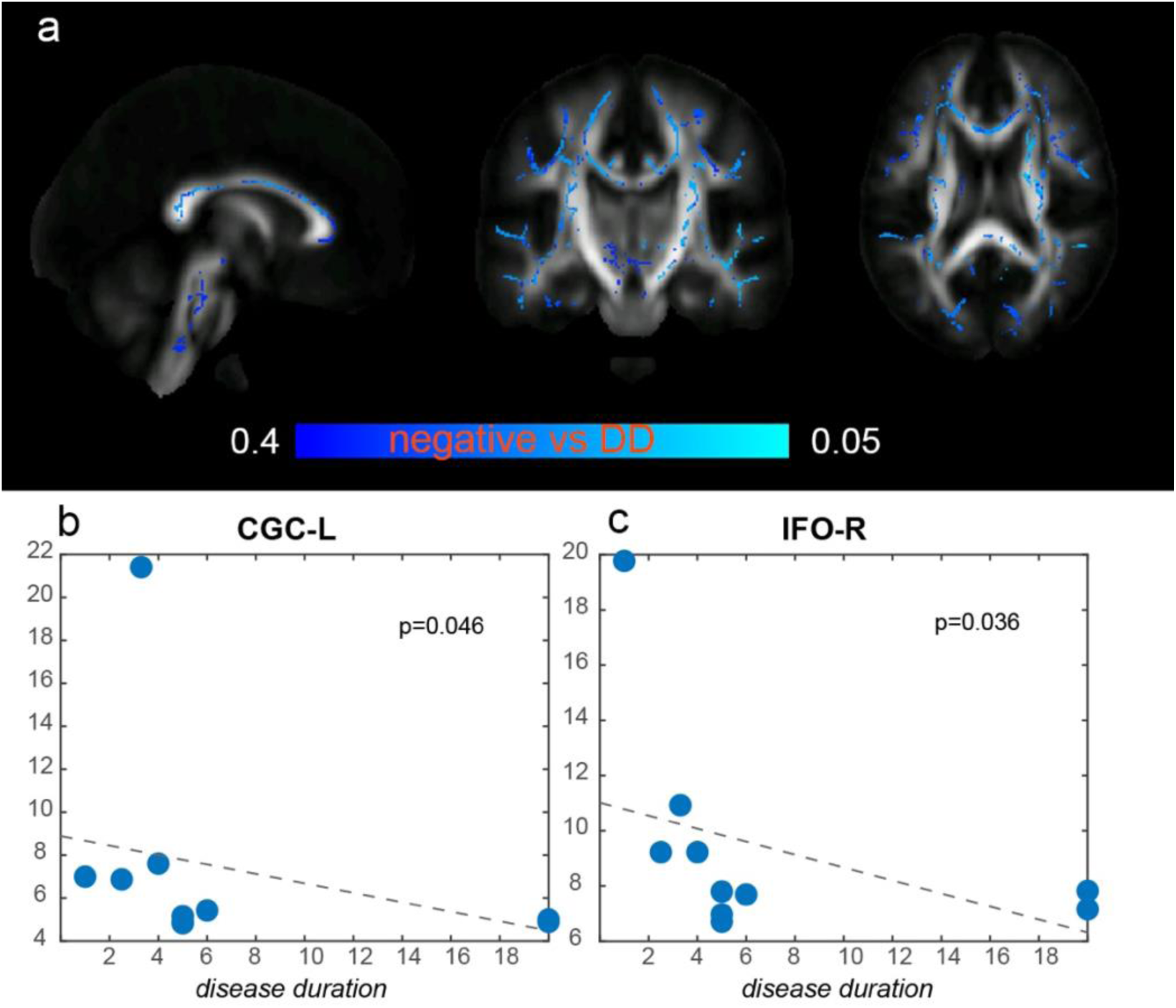
Negative association between axonal diameter and disease duration in TBSS (a) and in ROI-based analyses (b-c).

## Bibliography

Alexander DC, Hubbard PL, Hall MG, Moore EA, Ptito M, Parker GJM, Dyrby TB. 2010. Orientationally invariant indices of axon diameter and density from diffusion MRI. NeuroImage 52:1374–1389. doi:10.1016/j.neuroimage.2010.05.043

Alexander Leemans, Ben Jeurissen, Jan Sijbers, Derek K. Jones. n.d. ExploreDTI: a graphical toolbox for processing, analyzing, and visualizing diffusion MR data.

Assaf Y, Blumenfeld-Katzir T, Yovel Y, Basser PJ. 2008. Axcaliber: A method for measuring axon diameter distribution from diffusion MRI. Magn Reson Med 59:1347–1354. doi:10.1002/mrm.21577

Barazany D, Basser PJ, Assaf Y. 2009. In vivo measurement of axon diameter distribution in the corpus callosum of rat brain. Brain 132:1210–1220. doi:10.1093/brain/awp042

Bergers E, Bot JCJ, De Groot CJA, Polman CH, Lycklama a Nijeholt GJ, Castelijns JA, van der Valk P, Barkhof F. 2002. Axonal damage in the spinal cord of MS patients occurs largely independent of T2 MRI lesions. Neurology 59:1766–1771. doi:10.1212/01.WNL.0000036566.00866.26

Conforti L, Gilley J, Coleman MP. 2014. Wallerian degeneration: an emerging axon death pathway linking injury and disease. Nat Rev Neurosci 15:394–409. doi:10.1038/nrn3680

Constantinescu CS, Farooqi N, O’Brien K, Gran B. 2011. Experimental autoimmune encephalomyelitis (EAE) as a model for multiple sclerosis (MS): EAE as model for MS. British Journal of Pharmacology 164:1079–1106. doi:10.1111/j.1476-5381.2011.01302.x

Criste G, Trapp B, Dutta R. 2014. Axonal loss in multiple sclerosisHandbook of Clinical Neurology. Elsevier. pp. 101–113. doi:10.1016/B978-0-444-52001-2.00005-4

De Santis S, Drakesmith M, Bells S, Assaf Y, Jones DK. 2014. Why diffusion tensor MRI does well only some of the time: Variance and covariance of white matter tissue microstructure attributes in the living human brain. NeuroImage 89:35–44. doi:10.1016/j.neuroimage.2013.12.003

De Santis S, Granberg T, Ouellette R, Treaba CA, Herranz E, Fan Q, Mainero C, Toschi N. 2019a. Evidence of early microstructural white matter abnormalities in multiple sclerosis from multi-shell diffusion MRI. NeuroImage: Clinical 22:101699. doi:10.1016/j.nicl.2019.101699

De Santis S, Herranz E, Treaba CA, Barletta V, Mehndiratta A, Mainero C, Toschi N. 2019b. Whole brain in vivo axonal diameter mapping in multiple sclerosis2019 41st Annual International Conference of the IEEE Engineering in Medicine and Biology Society (EMBC). Presented at the 2019 41st Annual International Conference of the IEEE Engineering in Medicine & Biology Society (EMBC). Berlin, Germany: IEEE. pp. 204–207. doi:10.1109/EMBC.2019.8856433

Garcia-Hernandez R, Cerdán Cerdá A, Carpena AT, Drakesmith M, Koller K, Jones DK, Canals S, De Santis S. 2020. Mapping microglia and astrocytes activation in vivo using diffusion MRI (preprint). Neuroscience. doi:10.1101/2020.02.07.938910

Garcia-Hernandez R,Cerdán Cerdá A, Trouve Carpena A, Drakesmith M, Koller K, Jones DK, Canals S, De Santis S. 2022. Mapping microglia and astrocyte activation in vivo using diffusion MRI. Sci Adv 8:eabq2923. doi:10.1126/sciadv.abq2923

Haines JD, Inglese M, Casaccia P. 2011. Axonal Damage in Multiple Sclerosis: A XONAL D AMAGE IN MS. Mt Sinai J Med 78:231–243. doi:10.1002/msj.20246

Horowitz A, Barazany D, Tavor I, Bernstein M, Yovel G, Assaf Y. 2015a. In vivo correlation between axon diameter and conduction velocity in the human brain. Brain Struct Funct 220:1777–1788. doi:10.1007/s00429-014-0871-0

Horowitz A, Barazany D, Tavor I, Yovel G, Assaf Y. 2015b. Response to the comments on the paper by Horowitz et al. (2014). Brain Struct Funct 220:1791–1792. doi:10.1007/s00429-015-1031-x

Huang SY, Tobyne SM, Nummenmaa A, Witzel T, Wald LL, McNab JA, Klawiter EC. 2016. Characterization of Axonal Disease in Patients with Multiple Sclerosis Using High-Gradient-Diffusion MR Imaging. Radiology 280:244–251. doi:10.1148/radiol.2016151582

Inglese M, Bester M. 2010. Diffusion imaging in multiple sclerosis: research and clinical implications. NMR Biomed 23:865–872. doi:10.1002/nbm.1515

Jeurissen B, Leemans A, Tournier J-D, Jones DK, Sijbers J. 2013. Investigating the prevalence of complex fiber configurations in white matter tissue with diffusion magnetic resonance imaging: Prevalence of Multifiber Voxels in WM. Hum Brain Mapp 34:2747–2766. doi:10.1002/hbm.22099

Jones DK, Alexander DC, Bowtell R, Cercignani M, Dell’Acqua F, McHugh DJ, Miller KL, Palombo M, Parker GJM, Rudrapatna US, Tax CMW. 2018. Microstructural imaging of the human brain with a ‘super-scanner’: 10 key advantages of ultra-strong gradients for diffusion MRI. NeuroImage 182:8–38. doi:10.1016/j.neuroimage.2018.05.047

Keil B, Blau JN, Biber S, Hoecht P, Tountcheva V, Setsompop K, Triantafyllou C, Wald LL. 2013. A 64-channel 3T array coil for accelerated brain MRI. Magn Reson Med 70:248–258. doi:10.1002/mrm.24427

Klein A, Andersson J, Ardekani BA, Ashburner J, Avants B, Chiang M-C, Christensen GE, Collins DL, Gee J, Hellier P, Song JH, Jenkinson M, Lepage C, Rueckert D, Thompson P, Vercauteren T, Woods RP, Mann JJ, Parsey RV. 2009. Evaluation of 14 nonlinear deformation algorithms applied to human brain MRI registration. NeuroImage 46:786–802. doi:10.1016/j.neuroimage.2008.12.037

Kurtzke JF. 1983. Rating neurologic impairment in multiple sclerosis: An expanded disability status scale (EDSS). Neurology 33:1444–1444. doi:10.1212/WNL.33.11.1444

Lasič S, Nilsson M, Lätt J, Ståhlberg F, Topgaard D. 2011. Apparent exchange rate mapping with diffusion MRI. Magn Reson Med 66:356–365. doi:10.1002/mrm.22782

Loevner LA, Grossman RI, McGowan JC, Ramer KN, Cohen JA. 1995. Characterization of multiple sclerosis plaques with T1-weighted MR and quantitative magnetization transfer. AJNR Am J Neuroradiol 16:1473–1479.

Luchicchi A, Hart B, Frigerio I, Dam A, Perna L, Offerhaus HL, Stys PK, Schenk GJ, Geurts JJG. 2021. Axon-Myelin Unit Blistering as Early Event in MS Normal Appearing White Matter. Ann Neurol 89:711–725. doi:10.1002/ana.26014

Nikić I, Merkler D, Sorbara C, Brinkoetter M, Kreutzfeldt M, Bareyre FM, Brück W, Bishop D, Misgeld T, Kerschensteiner M. 2011. A reversible form of axon damage in experimental autoimmune encephalomyelitis and multiple sclerosis. Nat Med 17:495–499. doi:10.1038/nm.2324

Petzold A. 2005. Neurofilament phosphoforms: Surrogate markers for axonal injury, degeneration and loss. Journal of the Neurological Sciences 233:183–198. doi:10.1016/j.jns.2005.03.015

Polman CH, Reingold SC, Banwell B, Clanet M, Cohen JA, Filippi M, Fujihara K, Havrdova E, Hutchinson M, Kappos L, Lublin FD, Montalban X, O’Connor P, Sandberg-Wollheim M, Thompson AJ, Waubant E, Weinshenker B, Wolinsky JS. 2011. Diagnostic criteria for multiple sclerosis: 2010 Revisions to the McDonald criteria. Ann Neurol 69:292–302. doi:10.1002/ana.22366

Smith SM, Jenkinson M, Johansen-Berg H, Rueckert D, Nichols TE, Mackay CE, Watkins KE, Ciccarelli O, Cader MZ, Matthews PM, Behrens TEJ. 2006. Tract-based spatial statistics: Voxelwise analysis of multi-subject diffusion data. NeuroImage 31:1487–1505. doi:10.1016/j.neuroimage.2006.02.024

Tallantyre EC, Bø L, Al-Rawashdeh O, Owens T, Polman CH, Lowe JS, Evangelou N. 2010. Clinico-pathological evidence that axonal loss underlies disability in progressive multiple sclerosis. Mult Scler 16:406–411. doi:10.1177/1352458510364992

Torkildsen Ø, Brunborg LA, Myhr K-M, Bø L. 2008. The cuprizone model for demyelination. Acta Neurol Scand 117:72–76. doi:10.1111/j.1600-0404.2008.01036.x

van den Elskamp I, Lembcke J, Dattola V, Beckmann K, Pohl C, Hong W, Sandbrink R, Wagner K, Knol D, Uitdehaag B, Barkhof F. 2008. Persistent T1 hypointensity as an MRI marker for treatment efficacy in multiple sclerosis. Mult Scler 14:764–769. doi:10.1177/1352458507087842

van der Kouwe AJW, Benner T, Salat DH, Fischl B. 2008. Brain morphometry with multiecho MPRAGE. NeuroImage 40:559–569. doi:10.1016/j.neuroimage.2007.12.025

Van Waesberghe JHTM, Kamphorst W, De Groot CJA, Van Walderveen MAA, Castelijns JA, Ravid R, Lycklama Nijeholt GJ, Van Der Valk P, Polman CH, Thompson AJ, Barkhof F. 1999. Axonal loss in multiple sclerosis lesions: Magnetic resonance imaging insights into substrates of disability. Ann Neurol 46:747–754. doi:10.1002/1531-8249(199911)46:5<747::AID-ANA10>3.0.CO;2-4

Winkler AM, Ridgway GR, Webster MA, Smith SM, Nichols TE. 2014. Permutation inference for the general linear model. NeuroImage 92:381–397. doi:10.1016/j.neuroimage.2014.01.060

Zinkand WC, Moore WC, Thompson C, Salama AI, Patel J. 1992. Ibotenic acid mediates neurotoxicity and phosphoinositide hydrolysis by independent receptor mechanisms. Molecular and Chemical Neuropathology 16:1–10. doi:10.1007/BF03159956

